# Influence of Vocal Feedback on Emotions Provides Causal Evidence for the Self-Perception Theory

**DOI:** 10.1101/510867

**Authors:** Louise Goupil, Petter Johansson, Lars Hall, Jean-Julien Aucouturier

## Abstract

Emotional reactions are usually accompanied by vocalizations whose acoustic features are largely impacted by the physiological state of the body. While many theoretical frameworks emphasize the role played by the perception of bodily changes in the emergence of emotional feelings, few attempts have been made to assess the impact of vocal self-perception in this process. Here, we address this question by asking participants to deliberate out loud about how they would feel in various imaginary situations while we covertly manipulate their voices in order to make them sound emotional. Perceiving these artificial expressive cues in their own voice altered participants’ inferences about how they would feel. Crucially, this effect of vocal self-perception on felt emotions was abolished when participants detected our manipulation either explicitly or implicitly. Beyond demonstrating that vocal self-perception plays a role in the emergence of emotions, these results provide causal evidence for self-perception theories.

## Introduction

Psychologists have long hypothesized that emotions result from the perception of physiological changes that arise in the body in response to an environmental event, contrary to the intuition that bodily changes actually follow feelings (James, 1890). Yet, the hypothesis that the perception of bodily changes plays a causal role in the emergence of emotions has proven difficult to prove empirically.

Studies focusing on proprioception attempted to demonstrate that modifying an agent’s facial configuration (e.g., artificially inducing a smile) changes her emotional state (Strack, Martin, & Stepper, 1988). This approach however has led to non-conclusive results, with a recent paper reporting a failure to replicate this effect in a large number of independent laboratories (Wagenmakers et al., 2016). Other studies focused on interoception, showing for instance that cardiac cycle impacts emotional judgements about fear faces (Critchley & Garfinkel, 2017). Yet, since it focuses on the perception of external stimuli rather than felt emotions, this line of research does not allow firm conclusions neither. Overall, it is not obvious how definitive causal evidence might be obtained from studying interoceptive and proprioceptive signals, since it appears difficult to manipulate afferent signals from, e.g., the heart, without inducing a detection of the manipulation in the agent, nor modifying the functioning of the heart itself.

Here, we overcome this limitation by focusing on the voice. Like proprioceptive and interoceptive signals, the voice is deeply modified during emotional episodes: for instance, joy is associated with higher pitch, intensity, and energy in high frequencies, while sadness corresponds to the opposite pattern (Scherer, 2003). Contrary to other bodily signals however, we recently discovered that vocal signals can be captured and altered without individuals necessarily noticing the manipulation, or modifying their vocal production, leading to situations in which artificial emotional cues can be introduced, and perceived as self-originated (Aucouturier et al., 2016). Crucially, participants’ self-reports revealed that perceiving such artificial affective cues in their own voice altered their mood. One interpretation might be that participants merely perceive an affective stimulus in their proximate environment that shifts their emotional state in the same way that listening to pleasant music can lift your mood. Alternatively, here we hypothesize and test the possibility that the impact of vocal affective cues on emotions reflects veridical self-perception, in the sense that it only arises because changes in one’s own voice are perceived as reflecting physiological changes that are happening in one’s own body.

## Method

To test this hypothesis, we asked participants to deliberate out loud about how they would feel in various imaginary situations. While they spoke, we covertly manipulated subtle acoustic cues in their voices in order to make them sound emotional. We predicted that covert manipulation of affective cues in the voice should impact participants emotional reactions in the congruent direction (e.g., happy cues should induce positive reactions), but that this effect should disappear when speakers either i) detect our manipulation explicitly or ii) detect our manipulation implicitly, as evidenced through vocal compensation. Forward models of speech production state that a vocal signal matching internal predictions about the auditory consequences of planned motor articulations will be perceived as self-originated (Tian & Poeppel, 2015). We hypothesized that perceiving affective variations in such a ‘self-generated’ signal would impact emotional experiences. By contrast, implicit detections that the vocal signal is not self-generated, that can be evidenced through acoustic compensation (Jones & Munhall, 2000), should lead to a dismissal of the affective cues it contains. In sum, the emotional impact of artificial vocal cues should be limited to the cases where they are perceived as self-generated, and do not lead to vocal compensation.

### Participants

Fifty-five participants took part in the experiment. Given the design of the experiment we aimed to test groups of 18 participants (see procedure), and since pilot testing suggested that approximately 1/3 of the participants detected the manipulation explicitly, 1/3 implicitly, and 1/3 showed no detection at all, we tested three groups of 18 participants. Seven participants had to be excluded due to technical problems with recording or real-time vocal transformations (i.e., failure of the software, or communication between the software and the experimental interface coded in python), leaving N = 48 participants in the final sample (female = 29; age = 23 +/-3.24).

### Stimuli

36 emotional scenarios were adapted and translated in French from (Wilson-Mendenhall, Barrett, & Barsalou, 2013). To ensure that participants would experience a wide variety of emotions, we selected 9 scenario per valence (positive/negative) and arousal (high/low arousal) quadrants (as validated by (Wilson-Mendenhall et al., 2013). We verified in an independent group of 10 participants with equivalent demographic background as our main sample (4 females, age = 28.5 +/-4.48) that they were indeed perceived as such in our population.

### Voice transformation

Subtle emotional cues were artificially introduced in participants’ speech in real time using the voice transformation technique introduced in Rachman et al. (2018). The software, called DAVID, uses a selection of digital audio effects such as pitch (i.e., fundamental frequency) shifting and spectral filtering to simulate cues of emotional expression in running speech, with a very-low latency compatible with real-time vocal production. For the *happy* manipulation, we applied a positive pitch shift (+50 cents) as well as a spectral modification aiming to simulate the impact of smiling (notch filter at 2880 Hz, gain = 3, Q = 0.74) (Ponsot, Arias, & Aucouturier, 2018). For the sad manipulation, we applied a negative pitch shift (−70 cents) as well as a spectral modification aiming to attenuate the power in high frequencies, resulting in a darker sound (low shelf filter, 8000 Hz, gain = 0.25, Q = 1) (Aucouturier et al., 2016; Scherer, 2003).

### Procedure

Participants were fitted with closed headsets minimizing the contamination from environmental noise and their non-manipulated voice (Beyerdynamics – DT770), and their voice was recorded with a DPA 4088 headset and an RME UCX Fireface sound card, that allows for a roundtrip latency inferior to 20ms. An Apple MacBook Pro running PsychoPy (Peirce, 2007) was used to control stimulus presentation, recording responses, as well as communicate in real time with the voice transformation software (DAVID) to apply the various voice transformation depending on experimental conditions. Participants were asked to read each of the scenarii out loud, before vocally describing how they would feel in such situation. To ensure that detection occurred in only a subpart of our sample, we used a cover story: participants were told that the experiment aimed at uncovering why and how emotions arise in various imaginary situations. While they talked, the participant’s voice was covertly manipulated to make it sound *happy* (i.e., higher pitch, brighter spectrum), *sad* (i.e., lower pitch, darker spectrum) or remained unchanged (i.e., *neutral* condition).

In a previous study using a between subject design (i.e., each participant heard her/his voice manipulated in only one direction), and a very slow introduction of the vocal-feedback manipulation, we found both that most participants did not detect that their voice had been manipulated, and that none of the participants displayed acoustic compensation (Aucouturier et al., 2016). Thus, in the present experiment, to ensure that a substantial number of participants would detect the vocal-feedback manipulation either explicitly or implicitly (as indexed through acoustic compensation), we changed the direction of the vocal-feedback manipulation every 4 trials, resulting in 9 blocks of 4 *happy*, *sad* or *neutral* trials. Pilot testing revealed that in this within-subject design, the percentage of participants who detected the manipulation increased as opposed to a between subject design where only one effect was slowly applied to each participants’ voice (Aucouturier et al., 2016). Vocal feedback manipulation, block order and scenarii were pseudo-randomized across participants, such that across groups of 18 participants, each scenario appeared in the *happy*, *sad* or *neutral* condition for 1/3 of the participants.

After reading and describing how they felt (which took on average 79 +/-42 seconds), participants had to summarize how they would feel on: 1/ a continuous valence scale ranging from “very negative” to “very positive”; 2/ a continuous arousal scale ranging from “not arousing” at all to “very arousing”, and 3/ to report how sure they were that they would feel this way on a continuous confidence scale ranging from “not confident at all” to “very confident”.

Following the experiment, participants were debriefed with progressive questions, in order to assess whether they detected the vocal feedback manipulation or not. Depending on their responses, they were given a score of detection ranging from 1 to 6 by the experimenter, who at this stage remained blind to the participant’s behavior during the experiment (1 = "you intentionally manipulated my voice; 2 = “my voice was higher / lower sometimes, and it changed during the experiment”; 3 = “my voice sounded strange sometimes, and it changed during the experiment”; 4 = “my voice sounded strange, and it was not only because I am not used to hearing myself through headphones”; 5 = “my voice sounded strange, but it is probably because I am not used to hearing myself through headphones” ; 6 = “my voice was absolutely fine”). Participants also filled in a questionnaire in order to assess their level of alexithymia (Zimmermann, Quartier, Bernard, Salamin, & Maggiori, 2007) as well as social anhedonia (Gaillard, Gourion, & Llorca, 2013) (see Supplementary Figure S1). At the end of the experiment, they were informed of the true purpose of the study.

### Pre-processing and analysis

For each trial and each participant, we computed the fundamental frequency for both the non-modified (i.e., natural) and the modified (i.e., transformed) voice recordings, using the Praat software (Boersma, 2001). Pitch was when normalized at the individual level to allow statistical analysis across participants that had very different mean fundamental frequency (e.g., males and females). The analysis revealed that, although generally effective (e.g., see Figure 1E and 2E), our pitch transposition algorithm sometimes failed because of the low vocal or audio quality of the recordings. We excluded trials in which the acoustic analysis revealed a failure of the voice transformation algorithm, as indexed by the fact that the difference between the pitch of the modified and non-modified voice did not show the intended transposition (e.g., for the sad condition, we excluded trials if the pitch of the modified voice was not lower than the pitch of the non-modified voice). This pre-processing lead to excluding 7% of the data.

**Figure 1:**
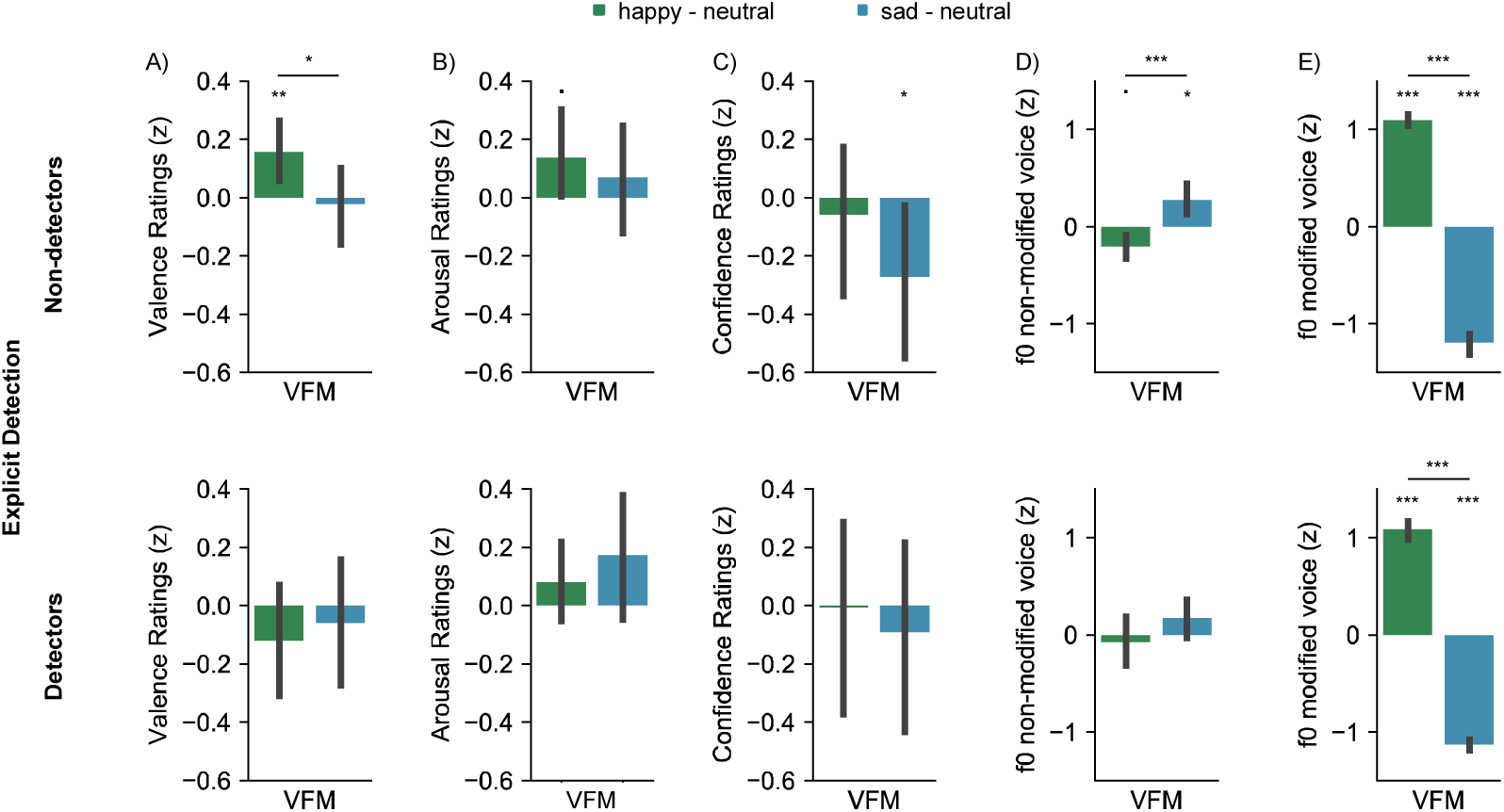
**(A)** Valence, **(B)** arousal, **(C)** confidence, **(D)** fundamental frequency of the non-modified voice (i.e. input of the manipulation) and **(E)** fundamental frequency of the modified voice (i.e. output of the manipulation) depending on the vocal-feedback manipulation (happy vs. sad) and explicit detection (top: participants who did not detected the manipulation; bottom: participants who reported the manipulation during the debriefing). Ratings were normalized with respect to neutral trials. Error bars represent confidence intervals - ^.^ represents p < 0.1; * p < 0.05, ** p < 0.01, *** p < 0.001.

For each participant, ratings in manipulated trials were normalized with respect to neutral trials, following this formula: *zX = (X – m_N)/sd_N*, were zX corresponds to normalized ratings, X to data samples, m_N to the average and sd_N to the standard deviation of the ratings given by this participant in neutral trials. Explicit detection was assessed from participants’ responses during the debrief: participants who detected that their voice had been manipulated (i.e., detection scores <= 4) constituted the group of explicit detectors, and participants who did not detect the manipulation at all (scores > 4) constituted the group of non-detectors.

Implicit detection was indexed by whether participants compensated for the acoustic manipulation in their vocal production. For each participant, acoustic compensation was assessed by testing whether the pitch extracted from non-modified voices (i.e., the vocal signal that was actually produced by the participant) differed from the participant’s average pitch. To do so, the pitch of non-modified voices was normalized, and tested against chance-level to assess their deviation from the participants usual range (pairwise comparisons between *happy* versus zero, and *sad* versus zero; significance threshold of < 0.1; note that similar results were obtained when normalizing with respect to neutral trials only). This analysis revealed that 50% (N = 24; 16 non-detectors) of the participants did not show any acoustic compensation; 16.7% (N = 8; 5 non-detectors) compensated in the happy condition (i.e., produced a lower pitch when their voice had been transposed upwards); 22.9% (N = 11; 7 non-detectors) compensated in the sad condition (i.e., produced a higher pitch when their voice had been transposed downward); 2% (N = 1; 1 non-detector) compensated in both directions; and 8.3% (N = 4; 2 non-detectors) of the participants displayed an outlying behavior (i.e., produced a higher pitch in the happy condition). Trials were classified as compensated if the participant showed a significant divergence in the pitch extracted for this condition (i.e., *sad* or *happy)*, and non-compensated if the pitch remained in the normal range for this condition. A rmANOVA confirmed that this allowed separating trials with substantial acoustic compensation at the level of the group (also see Figure 2D): there was an interaction between compensation and the vocal feedback manipulation (i.e., *compensated* vs. *non-compensated* trials; F(2,57) = 18 – p < 0.001) reflecting the fact that in *compensated* trials, the *sad* manipulation significantly increased the pitch of non-modified voices, as compared to both *neutral* (Tukey post-hoc comparisons with false discovery rate correction, p < .001) and *happy* trials (p < 0.001). The *happy* manipulation on the other hand decreased the pitch of non-modified voices as compared to *neutral* trials (p = 0.005). As expected, none of these comparisons were significant in non-compensated trials.

**Figure 2:**
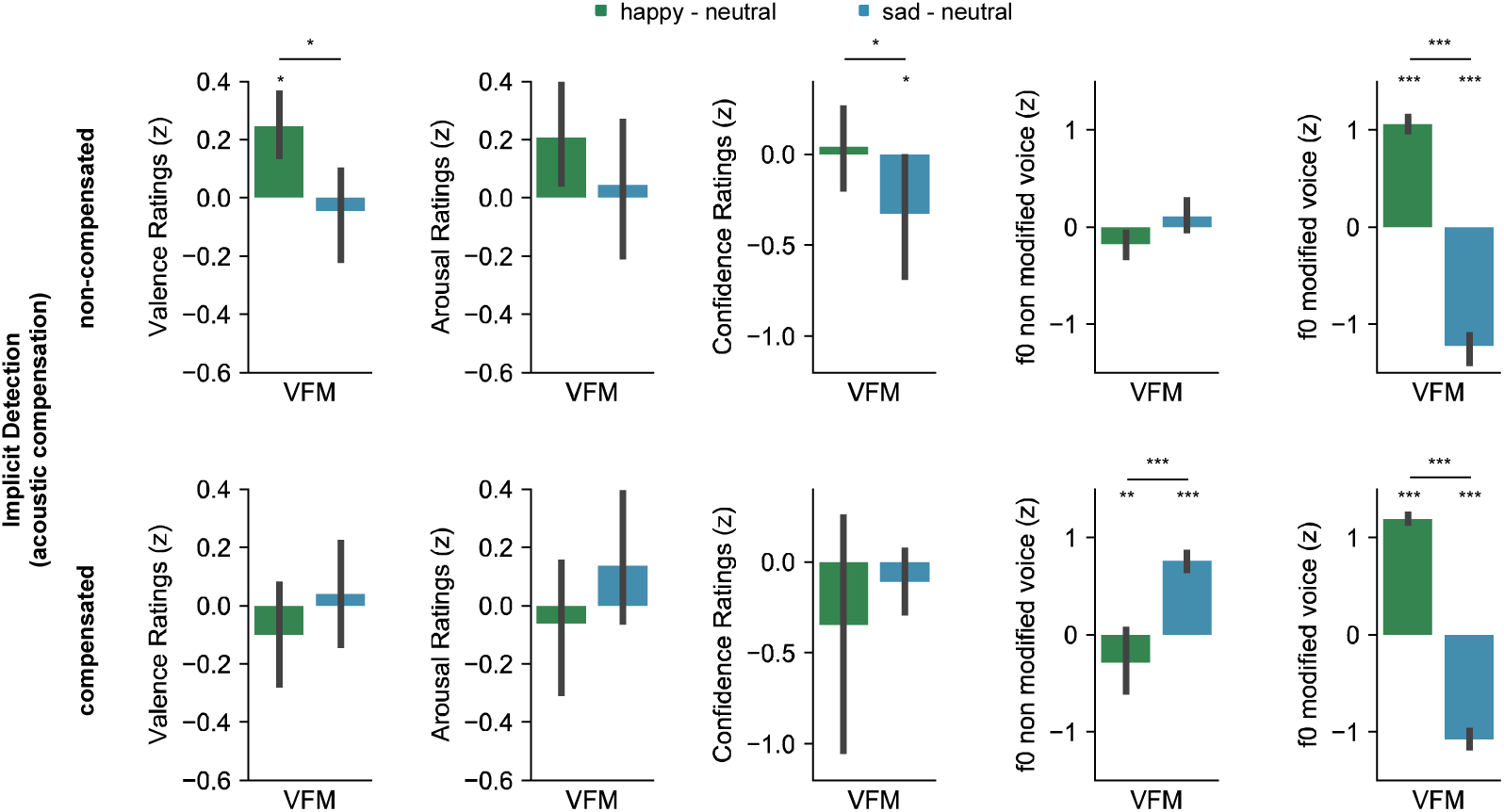
**(A)** Valence, **(B)** arousal, **(C)** confidence, **(D)** fundamental frequency of the non-modified voice (i.e. input of the manipulation) and **(E)** fundamental frequency of the modified voice (i.e. output of the manipulation) depending on condition (happy vs. sad) and implicit detection (i.e., acoustic compensation). Ratings were normalized with respect to neutral trials. Error bars represent confidence intervals - * p < 0.05, **p < 0.01, ***p < 0.001.

## Results

We first examined whether the Vocal Feedback Manipulation (hereafter VFM) had a main effect on the valence, arousal and confidence ratings provided by the participants. A repeated measure multi-variate analysis of variance (rMANOVA) revealed a significant effect of the VFM on our three main measures of valence, arousal and confidence ratings (F(2,94) = – Pillai = 0.17 – p = 0.01). Out of the 48 participants, 31 (65%) did not detect the manipulation explicitly (i.e., detection score of 5 or 6), and 17 (35%) participants detected that their voice had been manipulated (i.e., detection score <= 4). Consistent with our prediction, the main effect of the VFM on ratings was driven by the 31 participants that did not explicitly detect the vocal feedback manipulation (see Figure 1, compare top and bottom rows). For valence ratings, there was an interaction between the VFM and explicit detection (rmANOVA: F(2,92) = 3.66, p = .03, η*p*^2^ = .03) and a similar pattern was observed for arousal and confidence ratings, although the interaction did not reach significance (both F < 1 and η*p*^2^ < .01).

In non-detectors (N = 31), a rmMANOVA revealed a significant effect of the VFM on the three main measures (F(2,60) = 3.38 - Pillai = 0.29 - p < 0.01). Repeated measures analysis of variance revealed that there was a significant main effect of the VFM on both valence (F(2,60) = 5 - p < 0.01, η*p*^2^ = .08) and arousal ratings (F(2,60) = 3.16 - p < 0.05, η*p*^2^ = .04), and a marginal effect for confidence ratings (F(2,60) = 2.64 - p = 0.07, η*p*^2^ = .04) (see top row in Figure 1 A, B and C). Consistent with previous research (Aucouturier et al., 2016), the *happy* effect significantly increased valence ratings (M = 0.16 +/- 0.3; 95% CI [0.04, 0.27]; t(30) = 2.82, p < 0.01, d = 0.72), and marginally increased arousal ratings (M = 0.14 +/- 0.4 ; 95% CI [-0.02, 0.29]; t(30) = 1.75, p = 0.08, d = 0.45). This is consistent with our initial predictions (and the fact that in neutral trials we observed a relative increase of the fundamental frequency when participants reported positive emotions, see Figure S3), and suggests that the *happy* effect impacted participants emotional judgements in a congruent direction. The *sad* effect did not have a significant impact on valence (M = −0.02 +/- 0.35) nor arousal ratings (M = 0.06 +/- 0.5) overall, but it significantly decreased confidence ratings (M = −0.27 +/- 0.72; 95% CI = [-0.54, −001]; t(30) = −2.05, p < 0.05, d = 0.5, see Figure 1).

Based on previous research (Aucouturier et al., 2016), our initial prediction was that the *sad* effect should lead to a decrease in valence ratings, and no impact on arousal ratings. Several interpretations might explain the fact the *sad* manipulation did not impact valence ratings here. In particular, it appears that the impact of the *sad* VFM on emotional judgements was affected by contextual information (i.e., the baseline/intrinsic valence of the imaginary situation; see Figure S3 and discussion), while the happy effect was not. This reveals that affective cues can be interpreted differently depending on contextual information, and shows an interesting asymmetry between positive and negative situations.

Here it is important to note that although we labeled our effects “happy” and “sad”, it is likely that the cues that we introduced can actually be interpreted differently depending on the context. For instance, neutral states of low activation such as boredom have also been associated to lower pitch, while highly negative states of activation such as anger tend to be associated with higher pitch (Scherer, 2003). It might thus be that our “*sad”* effect leads to different inferences depending on the emotion that is afforded by a given trial’s scenario. The present experiment was not specifically designed to address this question, and follow-up experiments that carefully manipulate the emotional contents of the scenarii will be required to examine the fine interactions between contextual information and affective vocal cues during emotional appraisal.

Regarding confidence, although previous research has documented that epistemic uncertainty (i.e., doubting about one’s own state of knowledge) is generally signaled by a rising intonation and higher pitch (e.g., Jiang & Pell, 2017), the acoustic correlates of emotional uncertainty (i.e., doubting about one’s own emotional state) remained unclear. In the subset of our data were the voice remained non-manipulated (i.e., neutral trials), we found a lower pitch for trials where participants provided low confidence ratings (see Figure S3). Congruently, here we find that the sad effect decreases confidence in emotional judgements, thereby demonstrating for the first time that beyond impacting emotional judgements, vocal feedback also plays a role in “meta”-emotional judgements.

In any case, and crucially for our hypothesis, in detectors (N = 17), none of these effects were significant (all p-values > .2). Thus, consistent with our hypothesis, explicit detection of the vocal feedback manipulation abolished its impact on emotional self-reports.

We then turned to our marker of ‘implicit’ error detection in speech production, by examining acoustic compensation. We observed substantial acoustic compensation in the subgroup of 31 non-detectors, with a significant main effect of VFM on the fundamental frequency of the voice produced in manipulated trials (rmANOVA; F(2,60) = 8.4 – p < 0.001, η*p*^2^ = .2). At the level of the group, the *happy* manipulation lead the non-detectors to lower their voice as compared to neutral trials (M = -.20 +/- .36; 95% CI [-.36, -.001]; t(30) = −2, p < .05, d =.12), while the *sad* manipulation lead them to raise their voice (M = .40 +/- .8 ; 95% CI [.09, .5]; t(30) = 2.88, p < .01, d = 0.53, see Figure 1). At the individual level, we found that out of the 31 participants who failed to explicitly detect the VFM, 15 of them (31% of the total sample of N=48) showed substantial acoustic compensation for one or both of the VFM conditions. For each participant, trials were classified in two “implicit detection” conditions: *compensated* if the participant showed substantial acoustic compensation for that condition, and *non-compensated* if the participant did not show acoustic compensation for that condition (see methods).

Like explicit detections, implicit detections abolished the effect of the VFM (see Figure 2). Regarding valence ratings, a rmANOVA revealed a significant interaction between the VFM and implicit detection (F(2,57) = 3.55 – p < .04, η*p*^2^ = .05). No effect of the VFM was observable in the subset of the data corresponding to compensated trials (all p-values >

.3). In non-compensated trials however, the *happy* manipulation significantly increased valence ratings as compared to both *neutral* (Tukey post-hoc comparisons with false discovery rate correction, p < .04) and *sad* trials (p < .04). Although the interaction between VFM and compensation did not reach significance regarding arousal (p > .2, η*p*^2^ = .018) and confidence ratings (p > .2, η*p*^2^ = .024), post-hoc comparisons and restricted analysis suggest nonetheless a similar pattern (see Figure 2). In non-compensated trials, the *sad* manipulation significantly decreased confidence ratings as compared to both *neutral* (p < .05) and *happy* trials (p < .02), while in compensated trials, none of the effects were significant (all p-values > .2). Regarding arousal ratings, none of the effects reached significance (all p-values > .1). These results suggest that, as for explicit detection, implicit detection reflected in the presence of acoustic compensation somewhat blocked the impact of the VFM on emotional judgements.

Importantly, the disappearance of the effect in compensated trials was not trivially due to acoustic compensation somehow abolishing the vocal transformations: the pitch of *modified* voices in compensated trials still largely reflected the intended transposition in this subset of the data (see Figure 2, right panel). A rmANOVA showed that the effect of the VFM on the fundamental frequency of the voices remained highly significant in this subset of the data (F(2,57) = 15, p < 0.001, η*p*^2^ = .22), and there was no interaction between compensation and VFM (F < 1). In compensated trials, *happy* voices (M = 7.58 +/- 1.96) remained, as intended, significantly higher than neutral voices (Tukey post-hoc comparisons with false discovery rate correction p < 0.001) and *sad* voices (M = −3.71 +/- 1.7) significantly lower than neutral voices (p < .001) and *happy* voices (p < 0.001), right panel, Figure 2). Thus, the impact of the VFM was not abolished in compensators because acoustic compensation canceled our experimental manipulation. Rather, we suggest that acoustic compensation reflected an implicit detection that the affective cues currently perceived in the voice were not self-generated, leading to a dismissal of these cues during the construction of emotional judgement.

## Discussion

Perceiving artificial affective vocal cues impacted participants’ judgements about how they would feel in various imaginary situations. Specifically, introducing acoustic cues of happiness impacted emotions in a congruent direction, thereby replicating previous results (Aucouturier et al., 2016). Introducing affective cues of sadness also impacted emotional judgements, although this effect interacted with contextual cues. We also find that emotional uncertainty is associated with lower pitch, and that congruently, acoustic cues of sadness lower confidence ratings. Thus, second-order judgements about emotions can also be impacted by vocal signals. This is consistent with reports showing that confidence in perceptual judgements can be impacted by arousal (Allen et al., 2016), and calls for the development of embodied models of metacognition.

Crucially, the impact of vocal affective cues on emotions depended on these cues being perceived as self-generated: although most participants remained unaware of the manipulation, the substantial number of participants who detected the VFM showed no emotional effect. We also find that implicit detection that the vocal signal is not self-generated leads to a dismissal of the affective cues it contains. Extensive research has shown that perturbing speech can result in vocal compensation even when participants are not aware of the manipulation (Jones & Munhall, 2000), and that this phenomenon is related to self-identification (Tian & Poeppel, 2015). We demonstrate that, even in participants who do not consciously detect the manipulation, implicit detections indexed by acoustic compensations also abolish the influence of affective vocal cues on emotions. This is a stringent test of the hypothesis that only signals that are perceived as self-originated directly impact felt emotions.

The present study reveals that vocal self-perception plays a role in the emergence of emotions, and that this phenomenon is more complex than mere contagion, instead reflecting an inferential process integrating both the monitoring of physiological changes, and the interpretation of contextual information. This finding supports theories construing the perception and cognitive interpretation of bodily signals as central for the construction of idiosyncratic emotional experiences (Barrett, 2017; Damasio & Carvalho, 2013), but also suggest that, beyond interoception, exteroception of self-originated signals plays a substantial role in emotions.

## Supplementary Material

**Figure S1.**
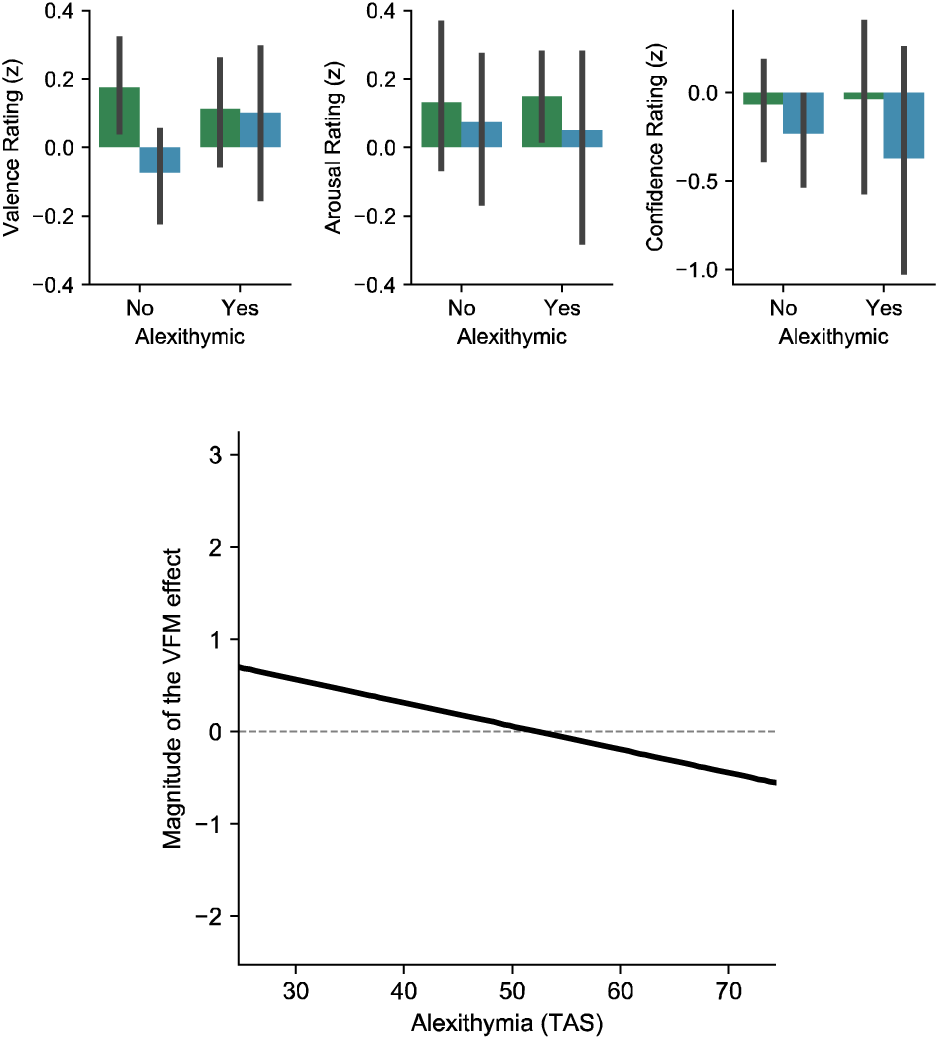
Relationship with Alexithymia. Several studies reported differences in the extent to which individuals are able to monitor their own interoceptive and bodily responses (Critchley & Garfinkel, 2017; Laird & Lacasse, 2014). Because a dysfunction in the perception of peripheral affective cues might impact the ability to monitor emotions, we examined whether self-reported levels of alexithymia impacted our effect. In our sample of 48 participants, 10 participants were alexithymic according to the TAS (i.e., TAS scores >= 61 correspond to high alexithymia, Zimmermann et al., 2007). They showed a reduced effect of the VFM (in this subgroup none of the main effects of the VFM were significant - all p-values > 0.05). In addition, there was an anti-correlation between TAS scores and a composite measure reflecting the VFM effect on emotional judgements (a sum of the differences between valence, arousal and confidence ratings in happy minus sad trials; values were reverse coded for confidence to allow comparison; rho = −0.36, p = 0.042). No such pattern was observed regarding social anhedonia. Although our sample size is not sufficient to allow firm conclusions here, it appears that the impact of vocal-feedback on emotions is reduced in alexithymia, and further research should seek to confirm this finding by specifically targeting patients suffering from alexithymia and control populations. Color code identical to main text (green: happy – neutral; blue: sad - neutral).

**Figure S2.**
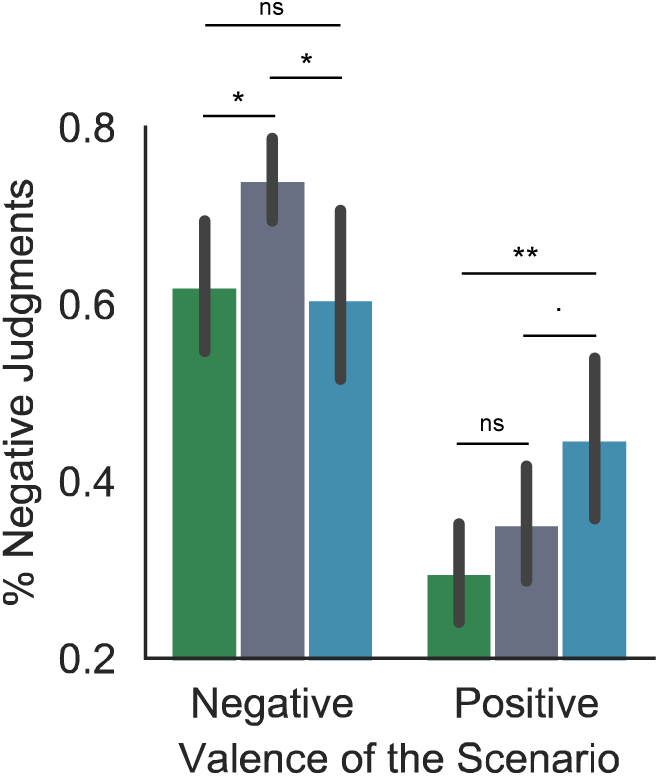
Integration of the VFM with the valence of the context. Splitting the data according to whether the scenario depicted a positive or a negative situation (as determined by a median split of valence ratings given by external participants, see methods), we observed that the *sad* effect was more dependent on the contextual cues contained in the scenario than the *happy* effect: the *sad* effect had opposite effects in positive vs. negative scenarii, but not the *happy* effect. Indeed, a rmANOVA on the percentage of negative responses revealed an interaction between the VFM and the valence of the scenario (F(2,34) = 4.42 - p < 0.02; only 18 participants had at least 1 data-point per cell for this analysis), and post-hoc tests showed that although for negative scenario the *sad* effect decreased the percentage of negative responses as compared to *neutral* trials (p < 0.02; Tukey post-hoc comparisons with false discovery rate correction), for positive scenario it marginally increased the percentage of negative responses as compared to *neutral* trials (p = 0.09), and significantly so as compared to happy trials (p < 0.01). ^.^ p < 0.1, * p < 0.05, ** p < 0.01, *** p < 0.001. Comparisons across the positive/negative dimensions are not shown for clarity, but all of them were highly significant (p < 0.01). Color code identical to main text (green: happy, grey: neutral; blue: sad)

**Figure S3.**
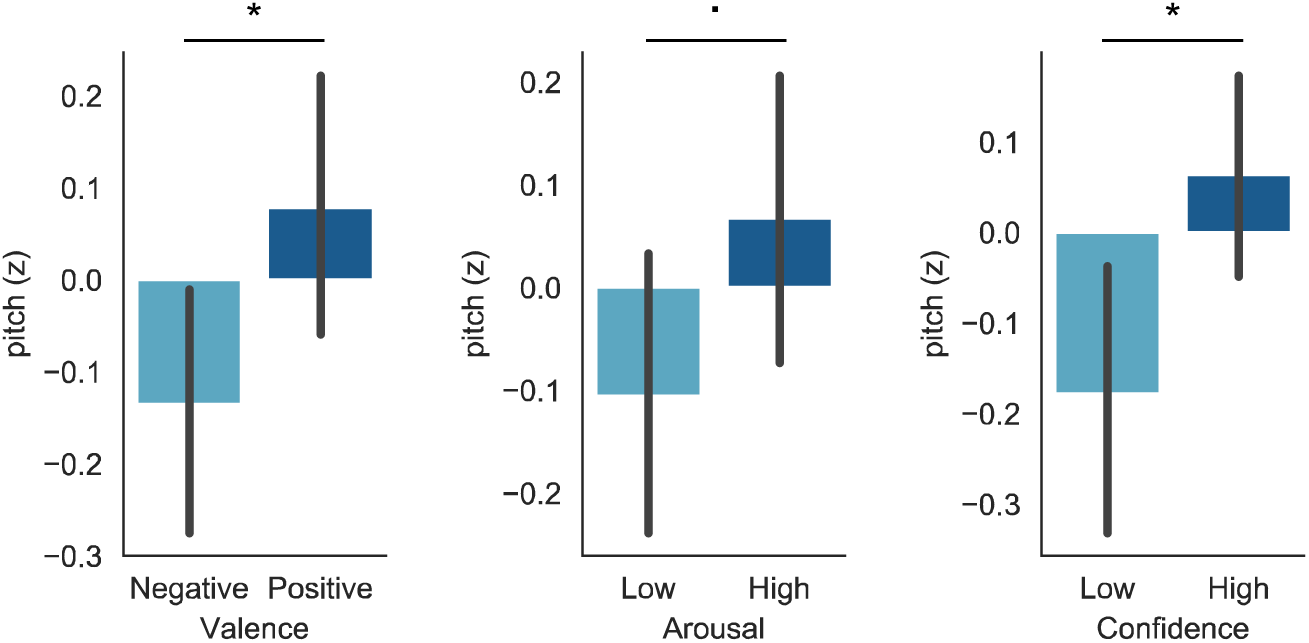
Impact of felt emotions on the voice. Fundamental frequency of the voices in neutral (i.e., non-modified) trials depending on whether participants reported feeling a negative or positive emotion (median split; left panel); high or low arousal (middle panel) and high or low confidence (right panel). Negative emotions were accompanied by a lower pitch as compared to positive emotions (t(47) = 2.17 - p < 0.04), and a similar pattern was observed for low confidence as compared to high confidence trials (t(46) = 2.8 - p < 0.01). Arousal marginally impacted pitch, with low arousal corresponding to lower pitch as compared to high arousal trials (t(47) = 1.96 - p = 0.056). This pattern is highly consistent with the impact of the vocal feedback on emotional judgements presented in Figure 1 and 2.

